# Comparative study of protein aggregation propensity and mutation tolerance between naked mole-rat and mouse

**DOI:** 10.1101/2021.08.20.457120

**Authors:** Savandara Besse, Raphaël Poujol, Julie G. Hussin

## Abstract

The molecular mechanisms of aging and life expectancy have been studied in model organisms with short lifespans. However, long-lived species may provide insights into successful strategies of healthy aging, potentially opening the door for novel therapeutic interventions in age-related diseases. Notably, naked mole-rats, the longest-lived rodent, present attenuated aging phenotypes in comparison to mice. Their resistance toward oxidative stress has been proposed as one hallmark of their healthy aging, suggesting their ability to maintain cell homeostasis, and specifically their protein homeostasis. To identify the general principles behind their protein homeostasis robustness, we compared the aggregation propensity and mutation tolerance of naked mole-rat and mouse orthologous proteins. Our analysis showed no proteome-wide differential effects in aggregation propensity and mutation tolerance between these species, but several subsets of proteins with a significant difference in aggregation propensity. We found an enrichment of proteins with higher aggregation propensity in naked mole-rat involved the inflammasome complex, and in nucleic acid binding. On the other hand, proteins with lower aggregation propensity in naked mole-rat have a significantly higher mutation tolerance compared to the rest of the proteins. Among them, we identified proteins known to be associated with neurodegenerative and age-related diseases. These findings highlight the intriguing hypothesis about the capacity of the naked mole-rat proteome to delay aging through its proteomic intrinsic architecture.

**Significance statement:** The molecular mechanisms behind naked mole-rat longevity are still poorly understood. Here, we address how the proteome architecture can help delay the onset of aging in naked mole-rat by studying properties that modulate protein aggregation. We identify ∼1,000 proteins with significant differences in aggregation propensity and mutation tolerance involved in processes known to be dysfunctional during aging. These findings highlight how evolutionary adaptations in protein aggregation in distinct biological processes could explain naked mole-rat longevity.

## Introduction

Understanding the mechanism of aging and life longevity is a major biological problem. The hallmarks of aging describe the dysfunction of several biological processes such as genomic instability, telomere attrition, loss of protein homeostasis (proteostasis), epigenetic alterations, mitochondrial dysfunction, cellular senescence, stem cell exhaustion, deregulated nutrient-sensing pathways, and altered intercellular communication (López-Otín et al. 2013). The aggravation of these hallmarks usually leads to an early manifestation of aging while their amelioration contributes to its delay and an increase of healthy lifespan. However, not all the hallmarks are fully supported yet by experimental interventions that succeeded in improving aging and extending lifespan. The genetics behind the hallmarks of aging have been identified through genetic perturbation studies in multiple model organisms such as yeast, nematodes, flies, and mice (reviewed in Singh et al. 2019; Taormina et al. 2019). These model organisms have been critical in our understanding of aging thanks to their short lifespan that aids tractable experimentation, relatively cheap maintenance, and possibilities for genetic manipulation. However, there is a need of studying organisms with longer lifespans, to better understand the mechanisms behind their longevity. Recent whole-genome sequencing efforts allowed the study of organisms with a longer lifespan. Cross-species “omics” studies of these long-lived species, such as unique transcriptomic, metabolic, and lipidomic profiles associated with long-lived species, highlighted molecular signatures that could be important to aging (reviewed in Ma and Gladyshev 2017; Tian et al. 2017). One notable example is the naked mole-rat, reported as the longest-lived rodent among those with a known maximum lifespan, which was recently used for studying healthy aging and longevity (Buffenstein and Ruby 2021). Indeed, this organism presents attenuated age-related changes, suggesting the presence of anti-aging mechanisms contributing to its longevity (Buffenstein, 2005). Several comparative studies between naked mole-rat and mice reported significant differences in the maintenance of protein homeostasis. Naked-mole rats show high oxidative damage levels from young ages (Andziak et al. 2006), but their ubiquitinylated proteins are maintained at lower levels at both young and old ages, suggesting a less accumulation of damaged and misfolded proteins during aging (Perez et al. 2009). The low levels of damaged and misfolded proteins could also be explained by their high proteasome activity (Rodriguez et al. 2012). Taken together, these observations emphasize the importance to study the general principles contributing to the robustness of protein homeostasis in the naked mole-rat. Nevertheless, these general principles have not been established at the proteome level. Thus, in this paper, we propose to identify the proteomic features that contribute to protein homeostasis maintenance.

In naked mole rats, several studies have previously studied the molecular key players of protein homeostasis and question their role toward rodent longevity. Proteostasis-centered theories of aging proposes that aging results from the decline of quality-control systems involved in protein synthesis, degradation, and chaperoning that normally contribute to protein turnover (Balch et al. 2008; Powers et al. 2009; Proctor and Lorimer 2011; Taylor and Dillin 2011). Proteostasis is essential for protein stability through the protection of their structures and functions against environmental perturbations. Impaired proteostasis leads to the appearance of phenotypic aging markers and age-related diseases such as Alzheimer’s and Parkinson’s diseases, known to be characterized by the accumulation of protein aggregates of specific proteins (Irvine et al. 2008; Powers et al. 2009; Hipp et al. 2019) Indeed, there is an increase in the expression of chaperones with higher proteasome and autophagy activities in naked mole-rat (Tian et al. 2017). From a system biology perspective, the maintenance of proteostasis is essential for delaying the onset or slowing down the process of aging (Koga et al. 2011)(ref). In addition, the protein aggregates are processed by quality control systems such as chaperones and protein degradation pathways (proteasome and autophagy) (Morimoto and Cuervo 2009).These mechanisms are robust in young individuals but tend to decline with age, leading to an increase of protein aggregates within the cell, thus participating in the dysfunction of multiple biological processes (Labbadia and Morimoto 2015). A recent study in *C. elegans* describes the proteostasis decline with age and observed an exponential increase of protein aggregates in old cells (Santra et al. 2019).

Our study focuses on intrinsic protein properties that could contribute to proteostasis maintenance by reducing the formation of protein aggregates. Causes of protein aggregation can arise from protein features and cell features. Protein aggregation propensity is a protein sequence feature that characterizes the ability of the protein to aggregate and is estimated based on the physicochemical properties of the amino acid sequence. Whether a sequence that has high aggregation propensity will in fact aggregate will need to account for cellular features. In the cell, cumulative damage through non-enzymatic post-translational modifications from reactions with metabolites or reactive oxygen species (Golubev et al. 2017), leads to protein instability, and subsequently to the formation of protein aggregates. Alternatively, the formation of protein aggregates could result from destabilizing mutations. The accumulation of somatic mutation burden has been proposed as a driver of aging (Vijg 2014). Several studies previously demonstrated the importance of mutation accumulation in the onset of aging and the reduction of lifespan (Lodato et al. 2018; Lee et al. 2019). However, it is still unclear whether the accumulation of mutation would contribute to the formation of protein aggregates. To tackle this question, we also propose to study “mutation tolerance” or the ability of proteins to tolerate the potential effects of mutations to increase their aggregation propensity. Here, we performed a comparative analysis on protein aggregation propensity and the mutation tolerance between the naked mole-rat and the mouse. From the study of these two properties, we aim to understand how they might contribute to explain the difference in lifespan between these two species.

First, we estimated their aggregation propensity for the whole-protein sequence, and the annotated domains of the proteins shared between the two species. We performed a random and exhaustive computational mutagenesis to estimate the mutation tolerance of these proteins. We found that although there is no global difference of aggregation propensity in the proteome shared between naked mole-rat and the mouse, we identified specific groups of proteins with significantly different in their aggregation propensity. This observation holds both at the level of individual domains and the level of entire protein sequences. By performing gene set enrichment analyses, we retrieve several biological processes, some of them were already reported to be potentially involved in the naked mole-rat longevity, notably processes associated with the immune system. We also highlight their inflammation’s versability, as we found proteins with high and low aggregation propensities from this process. We also report proteins, previously reported as involved in neurodegenerative diseases in human, that has not yet been considered as aging gene markers. Furthermore, these subsets of proteins have different distributions of mutation tolerance in the naked mole-rat, but not in the mouse, suggesting specific adaptations of these properties in the longest-lived rodent.

## Results

### Analysis of the orthologous proteome shared between naked mole-rat and mouse

To check the lifespan variability across rodents (Figure 1A), we collected maximum lifespan data available in the *AnAge* database (Tacutu et al. 2018) and retrieved information for 18 species. Furthermore, we extracted several metrics describing life-history traits such as body mass, basal metabolic rate, and female maturity available in *AnAge,* previously shown to be correlated with maximum lifespan in mammals (Fushan et al. 2015). On the reconstructed rodent phylogenetic tree, we observed that indeed the naked mole-rat is the longest-lived rodent (Ruby et al. 2018) and shares a common ancestor with other rodents living more than 12 years (Figure 1A, blue). This group is separated from a larger monophyletic group, which include a large cluster (Figure 1A, red) with rodents with a shorter maximum lifespan, less than 10 years, including mouse. The remaining two groups (Figure 1A, in green and orange) contain a low number of species with no clear tendency in their maximum lifespan. We plotted life-history traits metrics against the maximum lifespan (Figure 1B-D), confirming that naked mole-rat is an outlier from the rest of the rodents. These observations support the fact that the naked-mole rat is an appropriate organism to study aging because of its unexpectedly long lifespan among rodents, in contrast with mouse which shows up as a good representative for short-lived species.

**Figure 1:**
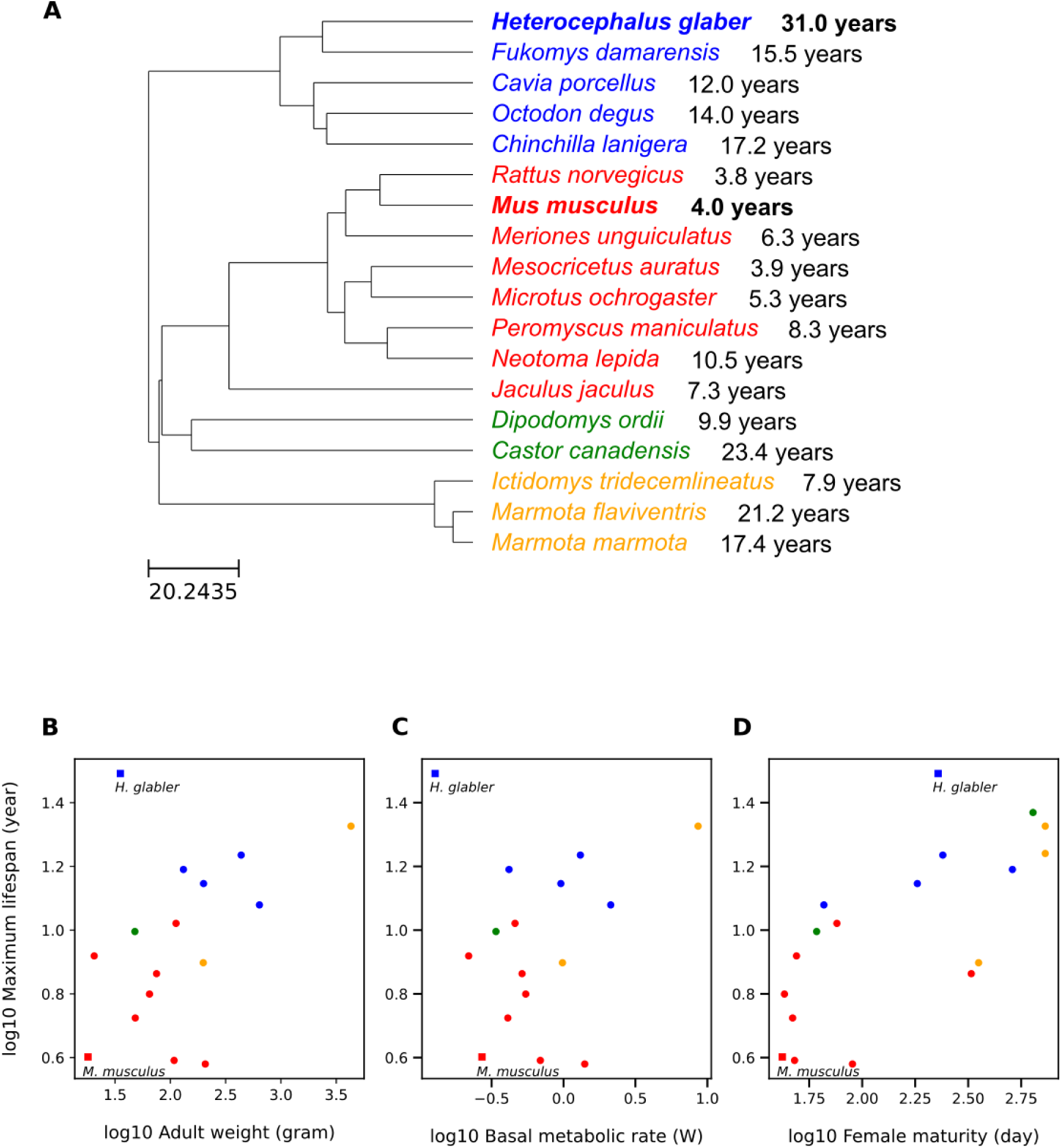
Maximum lifespan variation across rodents. (A) Phylogenetic distribution of rodent species with known maximum lifespan. The tree was generated with TimeTree using rodent species with known maximum lifespan. Four groups were colored according to their closest common ancestor. Mouse (*Mus musculus*) and naked mole-rat (*Heterocephalus glaber*), highlighted in bold, are the selected organisms for our comparative study. Rodent maximum lifespan compared to (B) adult weight, (C) basal metabolic rate, and (D) female maturity, for the rodents mentioned in A. Maximum lifespan, adult weight, female maturity, metabolic rate data are extracted from the *AnAge* database. All values were log10-transformed. Mouse and naked mole-rat are represented with a square shape.

To identify the general principles behind the naked mole rat’s longevity, we compared the orthologous proteome shared between naked mole-rat and mouse. The mouse has a well-curated and annotated genome and has also been extensively studied in the field of aging (Mitchell et al. 2015). Our comparative analysis between naked mole-rat and mouse focuses on 13,806 ortholog pairs collected with the orthologous mapping database *Inparanoid* (see Methods). We considered two properties among these orthologous proteins, specifically: 1) their aggregation propensity and their mutation tolerance, to see if they could partly explain the higher maintenance of protein homeostasis in naked mole-rat compared to the mouse. To study these properties within the two species, we estimated the aggregation propensity of the ortholog pairs using the software *Tango* (Fernandez-Escamilla et al. 2004) (see Methods), which scores the per-residue aggregation propensity of protein sequences. With this software, the property of protein aggregation propensity is accurately predicted on proteins with no transmembrane regions; therefore, we excluded the transmembrane proteins (see Methods), leaving a total of 9,522 ortholog protein pairs.

Since different regions of an ORF could have different folding properties, the aggregation propensity scores were also computed at the domain level. To do so, we retrieved 19,413 annotated domains available for 8,475 proteins (see Methods). Moreover, we looked more closely at a specific subset of proteins, the chaperone client proteins, which are the proteins interacting with chaperones in known protein-protein interaction networks. This subset is composed of 1,298 protein pairs (see Methods).

### Specific subsets of proteins display significant differences in aggregation propensity

The accumulation of protein aggregates is potentially toxic to cells (Stefani and Dobson, 2003) and result from the decline of protein homeostasis. Protein aggregation tends to increase with age and initiate amyloid-beta aggregation in *nematodes* and mice (Groh et al., 2017). Since such protein aggregates are found in specific tissues and cause age-related diseases such as Alzheimer’s and Parkinson’s diseases, we asked whether the systematic presence of protein aggregates within the cells could be correlated to the onset of aging. In the naked mole-rat, despite high levels of oxidation, they maintain low rates of ubiquitylated proteins (Perez et al. 2009), suggesting a reduced formation of protein aggregates. To identify if there is a proteome-wide difference in protein aggregation propensity between naked mole-rat and mouse, we first estimated the protein aggregation propensity on the ortholog proteins using Tango (see Methods). For a given protein sequence, this approach estimates the per-residue aggregation propensity scores based on their physicochemical properties with specific environmental parameters. With these scores, we computed two metrics, (1) an aggregation score for the whole-protein sequence and (2) an aggregation score for each annotated domain of the proteins (see Methods). We compared the aggregation scores between naked mole-rat and mouse, in the whole-protein sequence, and their domains (Figure 2).

**Figure 2:**
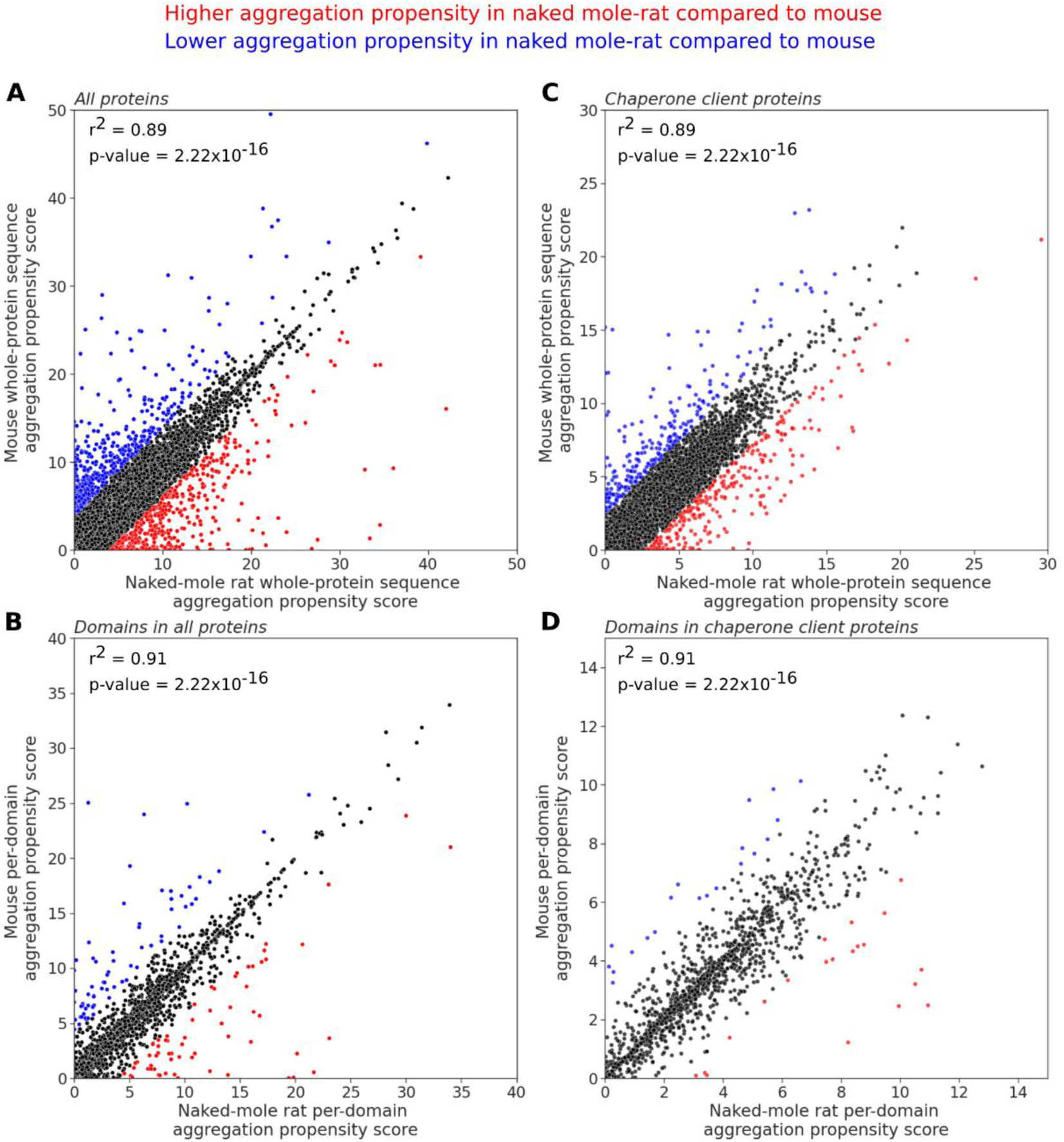
Study of aggregation propensity in naked mole-rat and mouse. Comparison of aggregation propensity scores in orthologous proteins from naked mole-rat and mouse. Each point represents an ortholog pair. Whole-protein sequence aggregation propensity scores (Agg_P_) (A) for the whole dataset (n=9,522), (B) for the subset of chaperone client proteins (n=1,298). Per-domain aggregation propensity scores (Agg _D_) (C) for the whole dataset (n=19,413 domains), (D) for the subset of chaperone client proteins (n=3,126 domains). See Methods for details on calculations Agg_D_ and Agg_P_. Pearson correlations coefficients (r^2^) between the naked mole-rat and mouse aggregation propensity scores are reported. Domains and proteins with a higher aggregation propensity in naked mole-rat compared to mouse are in red; and proteins with lower aggregation propensity are in blue.

Overall, whole-protein sequence propensity scores are low (Naked mole-rat Agg_P_=3.48 +/- 2.77, Mouse Agg_P_=3.37 +/- 2.73) and per-domain aggregation propensity scores have higher variance than whole-protein sequence (Naked mole-rat Agg _D_=3.79 +/- 4.60, Mouse Agg _D_=3.76 +/- 4.57). We observed a high correlation in aggregation propensity between naked-mole rat and mouse at the whole-protein sequence (r^2^=0.89, p-value=2×10^-16^) and the domain (r^2^=0.91, p-value= 2×10^-16^), indicating no proteome-wide global differences in aggregation propensity between these two species (Figure 2A,B). In parallel, we focused on the chaperone client proteins to see if these proteins have specific aggregation propensity and mutation tolerance compared to the rest of the proteins since they are interacting with the chaperones. Their whole-protein sequence and per-domain aggregation propensity scores are also low (Naked mole-rat Agg_P_=3.48 +/- 2.37, Mouse Agg_P_=3.37 +/- 2.31). We observed a high correlation in aggregation propensity, as in the all-proteins dataset, at the whole-protein sequence level (r^2^=0.89, p-value=2×10^-16^) and the domain level (r^2^=0.91, p-value=2×10^-16^) (Figure 2C,D), suggesting that chaperone client proteins do not differ in terms of aggregation propensity between these two species.

We computed differences of aggregation propensity (*ΔAgg*) to identify proteins differing significantly between the species. Altogether, we found 269 proteins (including 20 chaperone clients) with higher whole-protein sequence aggregation propensity (z-scores > 2, see Methods) in naked-mole rat compared to mouse, and 247 proteins (including 21 chaperone clients) with lower aggregation propensity (z-scores < -2). In proteins with annotated domains (n=8,475), we found 904 protein domains with a significant aggregation propensity score from 754 different proteins. Specifically, 452 protein domains (including 63 domains from chaperone clients) have higher aggregation propensity (z-scores > 2) in naked mole-rat compared to mouse, and 452 protein domains (including 70 domains from chaperone clients) have lower aggregation propensity (z-scores < -2). In total, in combining the whole-protein sequence and per-domain analyses, we identified 1,155 distinct proteins with differences in their aggregation propensity.

Additionally, we see no significant difference when comparing the distribution of *ΔAgg* z-scores from chaperone client proteins to proteome-wide values for the whole-protein sequence (p-value=0.72, t-test) and per-domain analyses (p-value=0.90, t-test). The proportion of proteins with a significant difference of aggregation propensity is similar in chaperone client proteins and the other proteins, indicating the chaperone client subset is not enriched in proteins with a significant difference of aggregation propensity between the naked mole-rat and mouse.

### Function of proteins with a significant difference of aggregation propensity

We investigated the over- and under-representation of specific Gene Ontology (GO) annotation terms associated of protein subsets with significantly high and low aggregation propensity in naked mole-rat (see Methods). We computed and sorted enrichment scores associated with each GO term (Figure 3). We found enriched or depleted groups having proteins with low aggregation propensity in naked mole-rat (in blue). These groups are associated with GO terms within Biological Process (Figure 3A) and Cellular Component (Figure 3B) categories. Depleted groups in *Biological Process* category are cell organization (5×10^-7^ < p-value < 7×10^-5^), regulation of different macromolecule biosynthesis (2×10^-8^ < p-value < 8×10^-5^) and regulation of gene expression (p-value=3×10^-5^). Proteins with significantly low aggregation propensity are under-represented in these processes. We found Ataxin-3 (ATX3) and Ataxin-10 (AT×10) proteins, associated with cell organization. These proteins are both responsible for different forms of spinocerebellar ataxia, a type of neurodegenerative disease. In contrast, enriched groups are related to immune response (p-value=8×10^-6^) and lipid metabolism (p-value: 4×10^-5^). We identified the amid ceramidase (ASAH1), an enzyme involved in lipid metabolism, and known to be associated with age-related diseases (Parveen et al. 2019). Depleted groups in *Cellular Component* category, are intracellular compartments, while enriched groups are membrane (p-value=5×10^-9^ & p-value=7×10^-5^) and extracellular components (Figure 3B), such as the extracellular matrix (p-value=1×10^-28^) and the cell surface (p-value=4×10^-7^). Notably, in these compartments, we found numerous metalloproteases from the matrixin family such as MMP3, MMP10, MMP13, MMP19 containing several hemopexin repeats; MMP7 and MMP25 with a peptidase M10 domain. These metalloproteases can degrade proteins from the extracellular matrix. Additionally, we noticed that proteins in the inflammasome complex (p-value=3×10^-6^) contain domains with significantly high aggregation propensity in naked mole-rat. Particularly, we identified the peptidase C14 domain of CASP-1 and CASP-12 from the caspase family, the NOD2-WH domain of NLRP-1A, NLRP-3, and NLRP-6, the functional domain of GSDMDC1, and the card domain of NLRC4. All these proteins are involved in inflammation.

**Figure 3:**
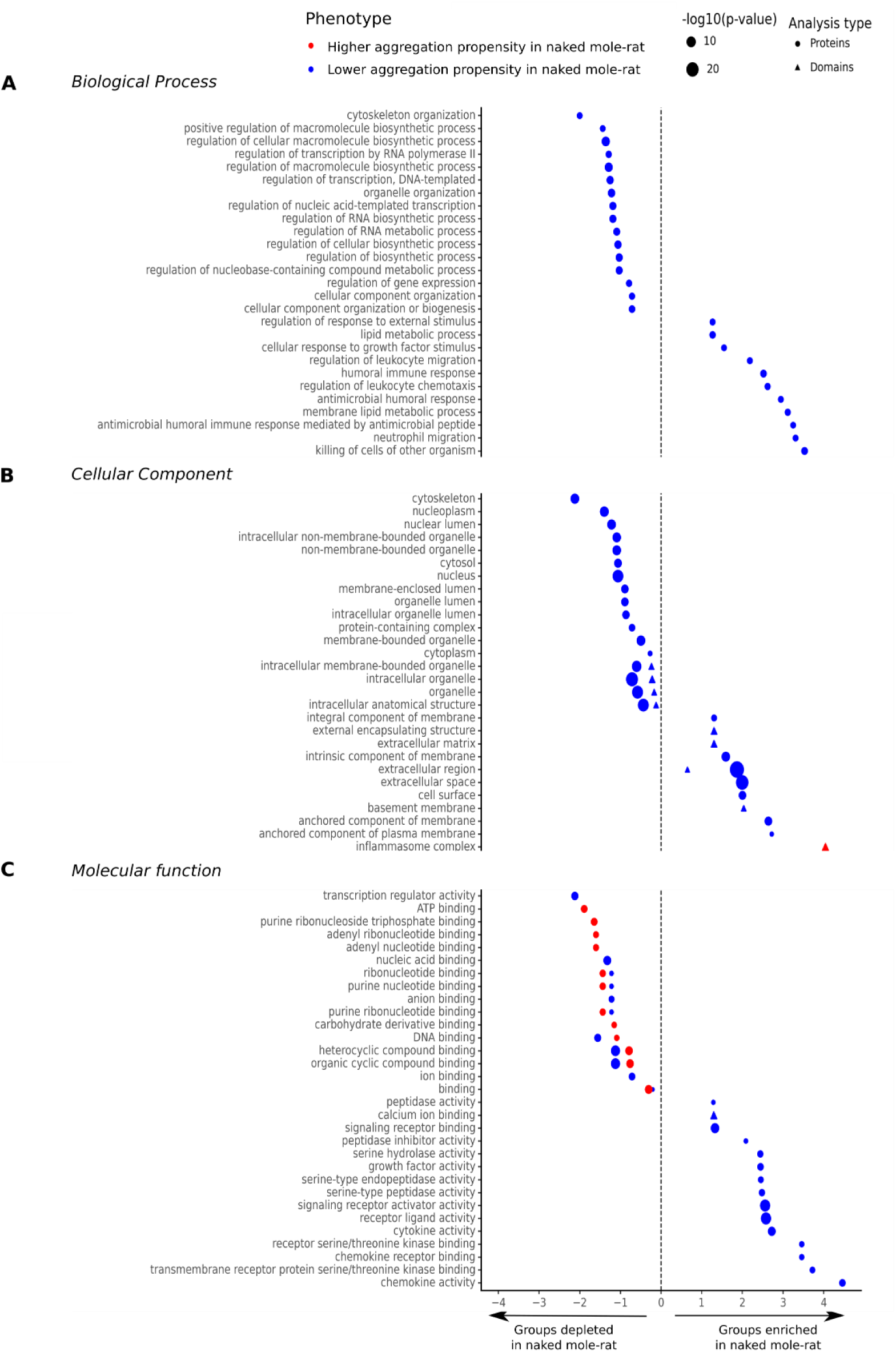
Significant Gene Ontology (GO) terms associated with domains and proteins with higher and lower aggregation propensity in naked mole rat. Log2 fold enrichment (FE) values indicate which GO terms are depleted (log2 FE < 0) or enriched (log2 FE > 0) in proteins (● shape) and domains (▲ shape) with a higher (in red) or a lower (in blue) aggregation propensity in naked mole-rat. The GO terms are grouped by categories: (A) Molecular Function, (B) Cellular Component, and (C) Biological Process. The size of the dots is proportional to their -log10 p-values. Only GO terms with at least 5 proteins and FDR < 0.05 are shown.

The lack of enriched GO terms for the subset of proteins with high aggregation propensity in naked mole-rat than in mouse across all GO categories suggest this may be a random group of proteins. However, in *Molecular Function* category (Figure 3C), we identified depleted groups in proteins with significantly high aggregation propensity (in red) related to ATP binding and its sub-categories (p-value=1×10^-5^). The associated proteins with these functions have a more conserved aggregation propensity than expected by chance. Interestingly, we found depleted groups containing both proteins with higher and lower aggregation propensity in naked mole-rat, related to various types of binding functions. This observation supports the fact that proteins with specific well-defined molecular functions are generally more structurally conserved across species and are therefore less likely to have significant differences of aggregation propensity between species. Nevertheless, only one group (calcium ion binding, p-value=3×10^-6^) contains proteins with differences of aggregation propensity from domains. The other enriched groups from *Molecular Function* category, contain proteins with different enzymatic activities (serine-type peptidase, p-value=3×10^-5^; serine hydrolase, p-value=3×10^-5^). Among them, we identified Chymotrypsin-C known to contribute to proteolysis, the breakdown of proteins as polypeptides. Finally, we found enriched groups of proteins associated with chemokine and cytokine activity (p-value=1×10^5^; p-value=1×10^-7^, respectively). We identified several members of the chemokine family, the immunoglobulin receptor IL-40, the interferon-alpha IFNA13, the Cerberus and Wnt-2b proteins from the Wnt pathway. All annotations of the proteins associated with specific GO terms are shown in Table S5 (Besse_et_al_SM.xlsx).

Furthermore, the distribution of the number of proteins per GO terms within each category (Figure S1) is similar for chaperone client proteins and the other proteins, except for the ones associated with immune response and extracellular components, (marked with an asterisk, corrected p-values < 0.05, chi-square test, Figure S1), indicating there are few or no proteins with lower aggregation in naked mole-rat that need chaperones to fold in these groups.

### Proteins with lower aggregation propensity in naked mole-rat better tolerate mutations

Finally, we explored the somatic mutation theory of aging through the study of mutation tolerance in naked mole-rat and mouse orthologous proteins. This theory hypothesizes that the accumulation of mutation is an essential player in the onset of aging (Kennedy et al. 2012) and influences longevity. We designed a large-scale *in silico* mutagenesis experiment by generating all possible 1-nucleotide mutations on gene sequences for 9,346 protein pairs (of length below 10,000 amino acids) and then estimating the aggregation propensity of these mutants (see Methods). The difference between the aggregation propensity from mutated sequenced and the aggregation propensity from the original sequences allows us to predict if a substitution would increase, maintain, or decrease this property. Assuming that proteins would preferably tolerate substitutions that do not significantly change their aggregation propensity, we derive a mutation tolerance score defined as a ratio of the number of substitutions with no change on the aggregation propensity divided by the total number of generated substitutions (see Methods). These values range from 0 to 1, representing weak to strong tolerance to substitutions, respectively.

This score allows us to study the relationship between whole-protein sequence aggregation propensity and mutation tolerance of orthologous proteins in the two rodents (Figure 4). There is a high correlation between mutation tolerance scores between naked mole-rat and the mouse (r^2^=0.89, p-value2×10^-16^), suggesting no global difference in mutation tolerance between their proteomes (Figure 4A). In both species, we observed a negative correlation between the sequence aggregation propensity and the mutation tolerance (r^2^=-0.58, p-value = 2×10^-16^), suggesting that proteins with a low aggregation propensity tend to be more resistant to substitutions (Figure 4B,C). Importantly, we identified subsets of proteins that have significant differences in mutation tolerance between the species (Figure 4A). We tested whether proteins with significant differences in mutation tolerance between the species have similar aggregation propensity than the rest of the dataset. The distribution of aggregation propensity for proteins with higher mutation tolerance compared to the distribution of other proteins is significantly different in both species (Figure 4B, Mouse p-value=1×10^-19^; Figure 4C Naked mole-rat p-value=2×10^-15^, Kolmogorov–Smirnov test), with proteins with higher mutation tolerance in a species having lower aggregation propensity compared to the rest of the proteins. This result implies that the proteins with low aggregation propensities better tolerate mutations which is not surprising, given that our mutation tolerance score itself is based on the whole-protein sequence aggregation propensity. We investigated the function of these proteins by performing an enrichment analysis as previously described, but no specific GO term was under- or over-represented in these subsets.

**Figure 4:**
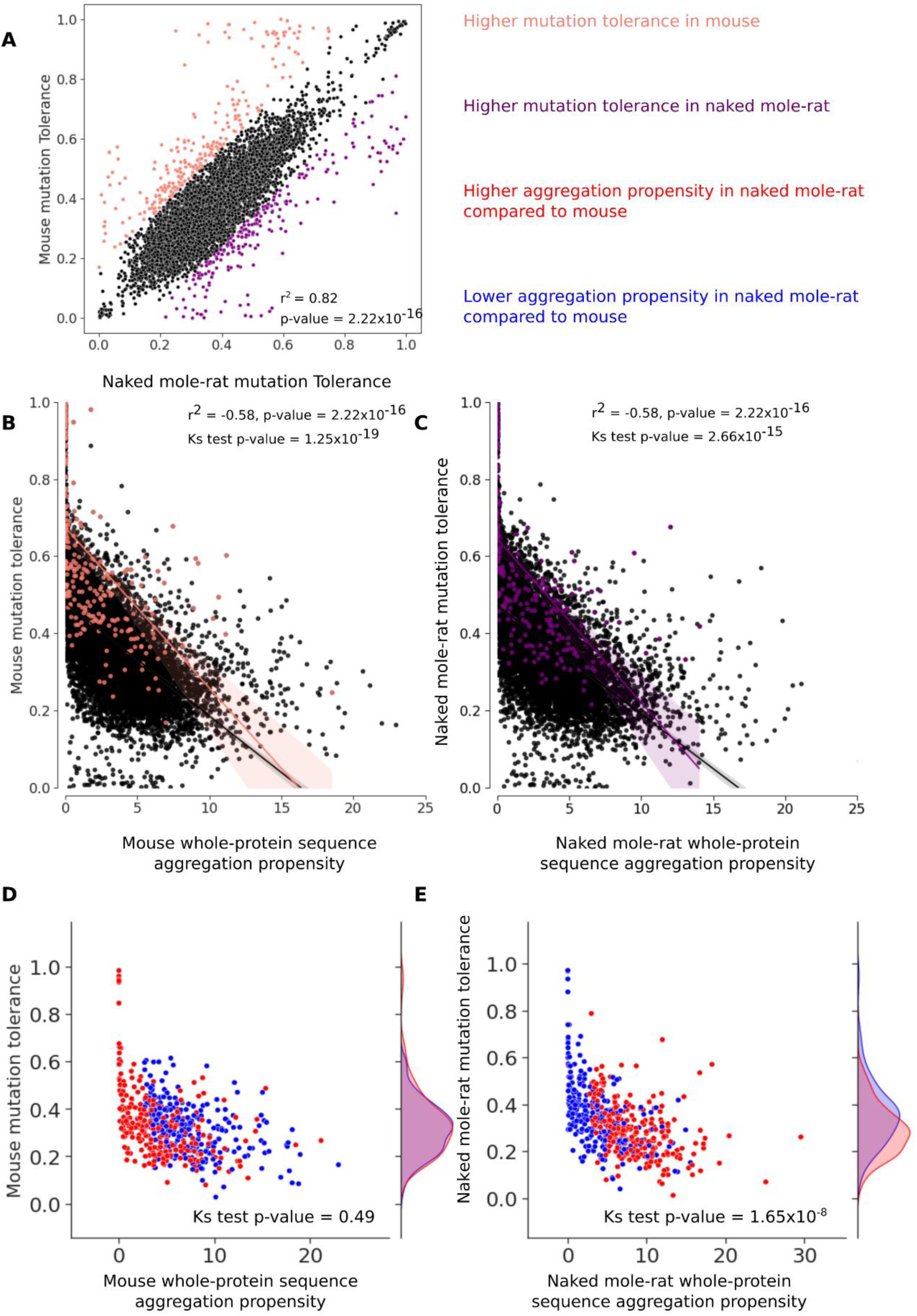
Study of mutation tolerance in naked mole-rat and mouse. (A) Comparison of mutation tolerance scores in orthologous proteins between naked mole-rat and mouse (n=9,346 proteins). Proteins in the naked mole-rat with higher mutation tolerance are in purple, the ones with lower mutation tolerance are in pink. Correlation between mutation tolerance and whole-protein sequence aggregation propensity scores in (B) mouse and (C) naked mole-rat. Protein pairs with significant differences in mutation tolerance are colored, using the color code from panel (A). Pearson correlation (r^2^) between mutation tolerance and aggregation propensity are reported in both organisms. Kolmogorov–Smirnov test is used to assess the difference of distribution between proteins with mutation tolerance scores similar in mouse and naked mole-rat, and the ones which are different. Scatterplots of mutation tolerance against whole-protein sequence aggregation propensity scores in (D) mouse and in (E) naked mole-rat, restricted to the subsets of proteins identified with significant difference of aggregation propensity (n=510 proteins). Proteins with higher aggregation in naked mole-rat compared to mouse are in red, proteins with lower aggregation are in blue. Kolmogorov–Smirnov (KS) test is used to assess differences in mutation tolerance distributions between the two subsets in each organism.

Lastly, we investigated the mutation tolerance scores of the proteins with higher and lower aggregation propensity in naked mole-rat compared to mouse. In the mouse (Figure 4E), the distributions of mutation tolerance scores between higher and lower aggregation propensity proteins are not significantly different (p-value=0.49, Kolmogorov–Smirnov test), indicating that the distributions of the mutation tolerance of the two subsets are similar. However, in naked mole-rat (Figure 4F), we find a significant difference in the mutation tolerance scores between higher and lower aggregation subsets (p-value=2×10^-8^, Kolmogorov–Smirnov test). In naked mole-rat, proteins with lower aggregation propensity better tolerate substitutions than proteins with higher aggregation propensity. These proteins are found in biological processes or pathways shown in Figure 3, which we will discuss as potential players towards naked mole-rat longevity.

## Discussion

Aggregation propensity and mutation tolerance are two intrinsic properties of proteins that could contribute to the better maintenance of protein homeostasis. In this study, we designed a computational strategy to estimate these properties at the scale of the whole-proteome in naked mole-rat and mouse using a comparative genomic framework. Among their orthologous proteome (n=9,522 proteins), we did not identify global difference in aggregation propensity, but about 1000 proteins showed significant differences, either from their domains or from their whole-protein sequences. In our analyses, we specifically study protein chaperone client proteins to determine whether this subset has differing intrinsic properties but did not find significant differences. Previous studies have shown that chaperone client proteins evolve slower and have a lower aggregation propensity compared to non-client proteins (Victor et al. 2020), but our study shows that these properties remain similar between naked mole-rat and mouse. As caveats, we inferred the naked mole-rat and mouse chaperone clients from human orthologs. It is therefore possible that the subset of proteins we defined as chaperone client proteins is highly incomplete or does not interact with chaperones in naked mole-rat and/or in mouse.

From the gene-enrichment analysis, we observed that the proteins of naked mole-rat with less aggregation propensity are over-represented mostly in the extracellular compartments, within several specific biological processes, related to immune response and lipid metabolism, and have functions associated with binding and protein degradation. The proteins with more aggregation propensity are not enriched in a particular biological process, except in the inflammasome complex, known to contain aggresomal complexes. Among the proteins we identified with significant differences in aggregation propensity, we identified several proteins previously known to be involved in neurodegenerative and age-related diseases. For instance, ATX3 is a poly-glutamine tract-containing protein, that contributes to cytoskeleton organization, and is known to be involved in protein inclusion bodies (Burnett and Pittman 2005). The accumulation of ATX3 in brain cells causes a proteostasis impairment that leads to the Machado-Joseph disease, or spinocerebellar ataxia-3 (Dantuma and Herzog 2020). Particularly, ATX3 is associated with double-stranded DNA binding. Previously, the study of ATX3-mutant in mouse brain cells showed an impairment of DNA repair efficiency, leading to the accumulation of DNA damage (Gao et al. 2015). ATA×10 was also identified here, which is associated with pentanucleotide disorder SCA10 (Bampi et al. 2017). The identification of lower aggregation propensity in these poly-glutamine proteins in naked mole-rat could contribute to resistance towards certain types of neurodegenerative diseases, leading to premature death (Dantuma and Herzog 2020). Moreover, we also identified proteins related to lipid metabolism with lower aggregation propensity in naked mole-rat, such as the acid ceramidase ASAH1. This proteins is involved in the intra-lysosomal ceramide homeostasis and is known to be associated with Alzheimer’s disease, cancer, and diabetes (Parveen et al. 2019). Furthermore, a recent study highlighted specific lipidic signatures in naked mole-rat that confer neuroprotective mechanisms against oxidative damage (Frankel et al. 2020). The lower aggregation propensity of the lipid metabolism proteins may contribute to protein stability and discharge of quality control systems of proteostasis.

Our study also highlighted the versatility of the aggregation propensity within inflammation pathways in naked mole-rats. Indeed, these rodents have a unique immune system able to better resist against bacterial infection. They have a unique myeloid cell subset that highly expressed genes for antimicrobial response (Hilton et al. 2019). Genes involved in the NOD-like receptor signaling pathway can activate pyroptosis, which is cell death after exposure to a bacterial infection. Interestingly, the NLRP-3 inflammasome pathway, which we found to have higher aggregation propensity at the level of protein domains in our study, is known to be regulated by the ubiquitin system. However, the exact molecular mechanisms of its non-canonical activation remain unclear (Lopez-Castejon 2020). The increase of domain aggregation propensity within proteins associated with the inflammasome complex could explain their affinity with the ubiquitin system. Moreover, during bacterial infection, naked mole-rat’s immune system is more frequently solicited than in the mouse (Cheng et al. 2017). In our study, we observed that proteins with chemokine and cytokine activity have significantly lower aggregation propensity. This suggest that the intrinsic properties of these naked mole-rat proteins adapt to be less prone to aggregate.

We also identified several metalloproteases having domains with lower aggregation propensity in naked mole-rat. Metalloproteases are known to degrade extracellular matrix proteins. Interestingly, the naked mole-rats highly produce the high-molecular-mass hyaluronan (Tian et al. 2013), a component of the extracellular matrix, known to have anti-inflammatory properties (Takasugi et al. 2020). These proteins might facilitate the hyaluronan turnover and balance the pro-inflammatory responses from the high activity of the inflammasome. Recently, two studies highlighted the importance of MMP13 as a therapeutic target for Alzheimer’s and Parkinson’s disease (Zhu et al. 2019; Sánchez and Maguire-Zeiss 2020). Tight regulation of inflammatory responses in naked mole-rat seems essential to maintain protein homeostasis, particularly during bacterial infection. Naked mole-rats are known to maintain proteasomal proteolytic activities in their late stages of life (Perez et al. 2009). These adaptations could indirectly promote healthy aging in naked mole-rat, increasing its maximum lifespan.

Mutation tolerance is another intrinsic property of proteins that could contribute to the maintenance of protein homeostasis. It indicates the ability of the protein to maintain its stability despite mutations. We used the difference of aggregation propensity between mutated and wild-type sequences to estimate whether a substitution event in the coding sequence would later drastically change or not the aggregation propensity of a protein. In the definition of our mutation tolerance score, synonymous substitutions favor protein stability and avoid the formation of protein aggregates. Despite no global differences in mutation tolerance between the two species’ proteomes, proteins with lower aggregation propensity in naked mole-rat better tolerate mutation compared to proteins with higher aggregation propensity. Such a difference is not seen in the mouse, which suggests these proteins in naked mole-rat have intrinsic properties that slow down the overload of the quality control systems of proteostasis, thus might contribute to its longevity.

Studying the diversity of lifespan within eukaryotes with comparative genomic approaches requires well-curated genome assemblies and reliable maximum lifespan measurements. In this study, we restricted our analysis to two species from the same taxonomic order, with a drastic difference of maximum lifespans, to identify the proteomic features explaining their difference of lifespan. Working with closed-related species helps to identify subsets of proteins specifically associated with biological processes related to longevity in the two species, without taking account of the complications arising from comparing from evolutionary-distant species Although these results could be specific to rodents, the pathways and genes identified in this study are known to be shared across eukaryotes. Therefore, our study is a first step towards a larger investigation of these properties across species. In addition to restricting the comparative analysis to only two species, our study has several limitations. First, we only focused on orthologous proteins shared between naked mole-rat and mouse, ignoring proteins unique to naked mole-rat which could also contribute to its extended longevity. Second, to predict the aggregation propensity of the proteins shared between naked mole-rat and mouse, we used the *Tango* software, which is a predictive approach that heavily relies on the physicochemical properties of the amino acid sequences and their likelihood to be involved in the formation of beta-sheets structures participating in functional folding. This approach performs well to predict the aggregation propensity of globular proteins (Linding et al. 2004), which resulted in the exclusion of transmembrane and membrane proteins from our analyses. Moreover, the aggregation propensity scores are predicted for a given set of environmental parameters and may not represent the dynamic range of aggregation propensity scores that the proteins could adopt in different tissues. Alternative bioinformatics methods to estimate aggregation propensity based on amino-acid sequences are implemented as webserver tools (Santos et al. 2020), incompatible with our high-throughput computational strategy for estimating mutation tolerance by generating billions of sequences that could only be processed in a timely manner using a command-line software. Therefore, *Tango* allowed us to build a systematic and highly efficient pipeline to estimate the aggregation propensity of ∼10,000 proteins in two different organisms, a large-scale experiment that is unfeasible to achieve *in vitro*. However, further molecular experiments will be necessary to validate the role of the identified less aggregation-prone proteins in naked mole-rat in the context of aging.

In conclusion, we investigated the peculiarity of naked mole-rat longevity by studying specific intrinsic properties of the proteome that influence the maintenance of proteostasis. Our study highlighted a trade-off in the regulation of inflammation responses in naked mole-rat, directly encoded in the amino acid composition of the proteins as it relates to its propensity to aggregation. We also identified several proteins with lower aggregation propensity compared to mouse that have been found to characterize neurodegenerative or age-related diseases in humans. Our findings propose the existence of a successful strategy encoded in the naked mole-rat proteome architecture to delay aging through better maintenance of protein homeostasis in the longest-lived rodent.

## Materials and Methods

### Definition of the orthologous dataset and subsets

Orthologous sequences are homologous sequences that share similarities from a speciation event. The orthologous amino acid (AA) sequences shared between naked mole-rat and mouse were retrieved using the *Inparanoid* algorithm (version 4.1) (Remm et al. 2001) with default parameters. As initial inputs, we use the naked mole-rat and mouse latest proteome assemblies, downloaded from Uniprot (https://www.uniprot.org/, accessed April 2019). The *Inparanoid* algorithm performs a reciprocal best-hit search to cluster the orthologous and in-paralog proteins, to identify the orthologous groups between the two species. For our analysis, each orthologous group was represented by a pair of proteins with the highest mutual best hit score, yielding 13,806 orthologous pairs. Mouse and naked mole-rat Uniprot protein identifiers are available in Supplementary Table S2 (Besse_et_al_SM.xlsx). To assess the quality of these orthologous pairs, we computed their local alignments with Matcher (Waterman and Eggert 1987; Huang and Miller 1991) and collected the percentage of similarity and the percentage of gaps within the pairwise alignments. Orthologous pairs with a percentage of similarity below 60% or a percentage of gaps above 20 % were removed, altogether keeping a total of 13,513 pairs.

For the estimation of aggregation propensity from Tango software (see below), we excluded transmembrane proteins. To identify the proteins with transmembrane regions to exclude, we first parsed mouse gene annotations available in the proteome FASTA file and defined the ones containing the keyword “transmembrane” as transmembrane proteins and excluded them. Additionally, we also predicted transmembrane regions in the remaining sequences with TMHMM (Krogh et al. 2001). All mouse and naked mole-rat proteins with at least one transmembrane region predicted were removed, restricting our analyses to 9,522 protein pairs. We also collected their associated protein-coding nucleotide sequences for our computational large-scale mutagenesis analysis (see below). Moreover, we identified a specific subset, containing all the proteins known to interact with chaperone proteins. For this specific dataset, we used the human chaperone client proteins (annotated with their ENSEMBL identifiers) from a recent study (Victor et al. 2020) to infer the mouse chaperone clients. The human ENSEMBL identifiers were converted to their corresponding Uniprot identifiers for mapping them towards the mouse Uniprot ortholog identifiers. Similarly, we then mapped the mouse Uniprot identifiers to the naked-mole rat ortholog identifiers. This specific subset of orthologs is composed of 1,298 protein pairs.

### Identification of protein domains in naked mole-rat and mouse

To obtain mouse and naked-mole rat domain definitions, we first collected mouse domain information from the Pfam database (http://pfam.xfam.org/, version 33.1). Within a given protein, we considered any peptide as a functional domain when their entire sequence matched domain annotations, corresponding to the start and end positions in PFAM protein alignments. For the naked-mole rat, the domain definitions were inferred using the reciprocal best hit method where the mouse annotated domains are used as reference. We collected a total number of 19,413 annotated domains available for 8,475 protein pairs, representing 89% of our initial dataset.

### Phylogenetic tree and data related to longevity

The evolutionary distances between rodent species were determined using *TimeTree* (Kumar et al. 2017), through the available webserver. This method retrieved all existing phylogenetic trees for the given species and provided the concatenation of these trees to determine the median time when species diverged. These phylogenetic trees were built based on gene alignments. The available information on maximum lifespan, adult weight, female maturity, and metabolic rate for rodent species was retrieved from the *AnAge* database (Tacutu et al. 2018, build 14) and are given in Table S1 (Besse_et_al_SM.xlsx)

### Computation of aggregation scores

To predict the propensity of proteins to aggregate, we used the Tango software (Fernandez-Escamilla et al. 2004). Tango assigns per-residue aggregation propensity scores based on the amino acid physicochemical properties. For each orthologous protein pair, we computed the per-residue aggregation score with Tango for each sequence independently and then calculated their whole-protein sequence aggregation and domain aggregation. Per-domain aggregation score is defined as the sum of the per-residue aggregation propensity score for a defined functional domain divided by the domain length (*Agg _D_*, Equation 1). The whole-protein sequence aggregation propensity score is defined as the sum of per-residue aggregation propensity scores for the entire sequence divided by the protein length (*Agg _P_*, Equation 2).

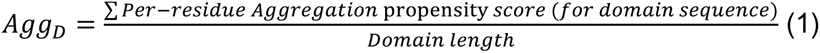

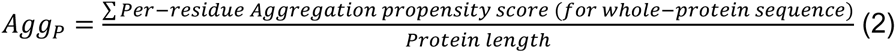

### Identification of proteins with significant difference of aggregation propensity

To compare mouse and naked-mole rat protein aggregation propensity scores, we computed their difference at the domain (*ΔAgg_D_*, Equation 3) and the whole-protein sequence (*ΔAgg_P_*, Equation 4) levels with the following formulas:

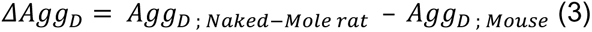

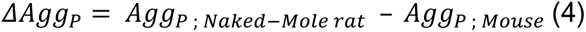

The difference of aggregation propensity scores was normalized to obtain z-scores. Proteins with z-scores exceeding 2 times the standard deviation are considered significantly different from each other. Both for whole-sequence and domain aggregation propensity analyses, two groups were defined as: (i) proteins with *ΔAgg* z-scores > 2 being considered to have a higher aggregation in naked-mole rat compared to mouse; (ii) proteins with *ΔAgg* z-scores < -2 being considered to have a lower aggregation in naked-mole rat compared to mouse.

### Functional enrichment analyses

With the previously identified subsets of proteins, we investigated in which cellular components, molecular functions, and biological processes from GO annotations, these proteins are over or under-represented. To do so, we used hypergeometric tests implemented on the Panther database (Mi et al. 2019). As the protein annotations for naked mole-rat were not proposed in the database, we used the annotations from the mouse, assuming the naked mole-rat proteins have similar annotations to their mouse orthologs. The subsets from the domain analysis were compared to the set of proteins with annotated domains within the shared proteome (n=8,475). The subsets from the whole-protein sequence analyses were compared to all the proteins of the shared proteome (n=9,522). Raw p-values of Fisher’s exact tests were computed to identify the gene ontologies significantly over- or under-represented for each subset, corrected by a False Discovery Rate (FDR). Only GO terms associated with at least 5 proteins are shown in Figure 3. The entire list of GO terms with FDR < 0.05 and, for the domain and the whole-protein sequence analyses, are available in Table S3 and S4, respectively (Besse_et_al_SM.xlsx). The list of proteins within the groups and their annotations are available in Table S5 (Besse_et_al_SM.xlsx).

To identify which GO terms where the chaperone client proteins are differently distributed compared to the rest of the proteins, we computed chi-square tests, corrected by a Benjamini/Hochberg FDR.

### Quantification of protein mutation tolerance

We designed a large-scale *in silico* mutagenesis experiment to estimate the mutation tolerance of the proteins shared between naked mole-rat and mouse. Specifically, the mutation tolerance score is a ratio from 0 to 1 that quantifies the ability of a protein to tolerate mutations. We mutated one nucleotide at a time within the DNA sequence to all 3 other possible nucleotide mutations (self-substitution is excluded). For example, for a coding sequence of X nucleotides, we would generate X^3^ possible substitutions that would engender X^3^ mutated sequences. All these DNA sequences are then translated into amino acid sequences. We kept only non-redundant protein sequences (resulting from non-synonymous changes), different from the wild-type sequence (WT), for predicting their protein aggregation propensity using Tango, as described in the section *Computation of aggregation scores*. Whole-protein sequence aggregation scores for mutated (MT) sequence were then computed and are used to calculate the difference of aggregation propensity (Mutational Agg _P_ - Equation 5) between MT and WT sequences:

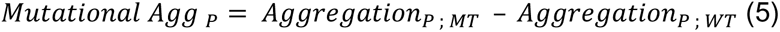

As Mutational Agg _P_ scores are normally distributed, we set a threshold at 2 times the standard deviation to distinct proteins with a significant change in their aggregation propensity score. We defined 3 categories of proteins, according to their change in aggregation propensity

1. Mutational Agg_P_>2: Increase in aggregation propensity of the mutated sequence
2. Mutational Agg_P_=0: No change in aggregation propensity of the mutated sequence
3. Mutational Agg_P_<-2: Decrease in aggregation propensity of the mutated sequence

For a given protein, these scores were used to define their mutation tolerance. It calculated the ratio of the number of mutations with no impact on protein aggregation normalized by the number of all possible mutations (Mutation tolerance, Equation 6).

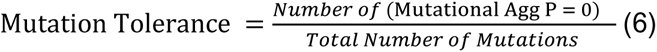

For identifying proteins with a significant difference in their mutation tolerance, we calculated the difference of mutation tolerance between naked mole-rat and mouse (ΔMutTol).

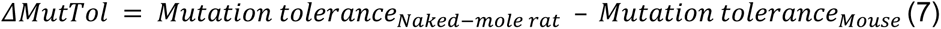

All the ΔMutTol scores were normalized to obtain ΔMutTol z-scores. Proteins with ΔMutTol z-scores exceeding 2 times the standard deviation are considered significantly different from each other: (1) proteins with a ΔMutTol z-score > 2 are considered to have higher mutation tolerance in naked-mole rat compared to mouse, (2) proteins with a ΔMutTol z-score < -2 are considered to have a lower mutation tolerance in naked-mole rat compared to mouse. This analysis was initially performed on 9,522 protein pairs. However, 176 proteins (mostly proteins with more than 10,000 amino acids) were removed as the calculation of their mutation tolerance score was too computationally expensive, thus, reducing the dataset to 9,346 protein pairs.

### Figure generation and statistical analysis

The different plots were generated with Python graphic libraries, Matplotlib (version 3.2.1), Seaborn (version 0.10.0), and Plotnine (version 0.8.0). All statistical analyses were performed using the Scipy stats module (version 1.6.2), unless specified otherwise. The FDR correction were computed with the statsmodels module (0.12.2), unless specified otherwise. Significance thresholds for p-values and FDR were set at 0.05. Statistical tests and p-values are reported in the figure legends can be found as outputs of the Python3 scripts that generate the figures.

## Availability of Data and Materials

The processed data and code used to generate the figures are available in the following Github repository: https://github.com/ladyson1806/NKR_lifespan. We also provide the different Python3 scripts and notebooks used to collect and pre-process the initial dataset, as well as the code that generates the different scores.

## Acknowledgments

This work was supported by funds from the Department of Biochemistry and Molecular Medicine of Université de Montréal and through the access to computational resources provided by Calcul Québec to JGH. We thank Sebastian Pechmann for his supervision on the preliminary analyses. We are grateful to Adrian Serohijos for his mentoring support and the fruitful discussions throughout the project. Finally, we thank all the members of the Hussin lab for their constructive comments and feedback on the figures for the manuscript. JGH is a Fonds de la Recherche du Québec en Santé (FRQS) Junior 1 Scholar, funded by the Institute for Data Valorization (IVADO).

