## Supplementary Materials for "Comparative study of protein aggregation propensity and mutation tolerance between naked mole-rat and mouse"

These Supplementary Materials contain the following:

- Supplementary Figure S1
- Legends for Supplementary Tables S1 through S5

### Supplementary Figure S1

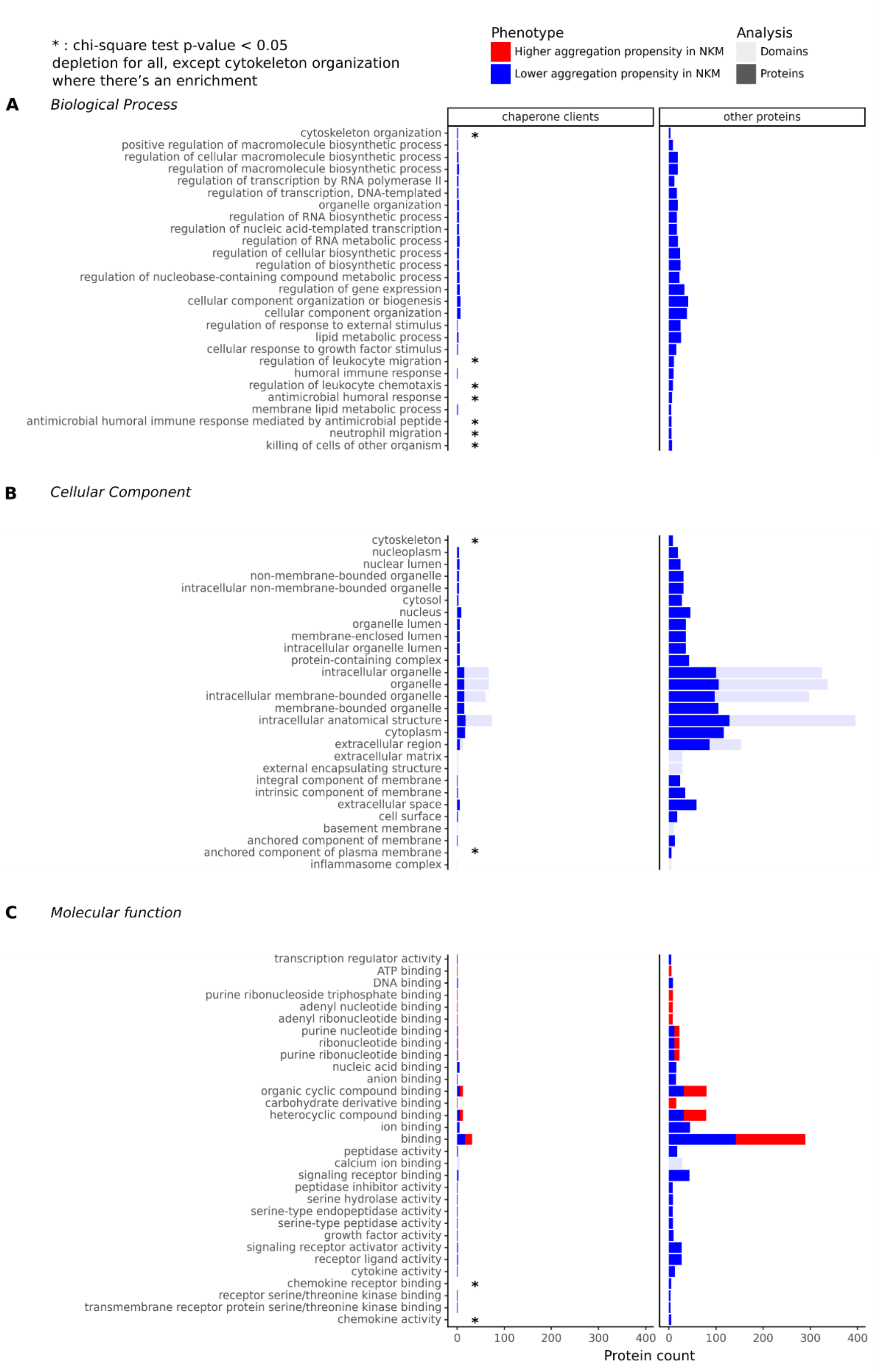

### Supplemental Figure legends

**Supplementary Figure S1: Protein count per GO terms associated to proteins and domains with significant difference of aggregation propensity**

This graph shows the number of proteins within each GO term. Redundant protein identifiers were not removed. (A) shows only chaperone client proteins and (B) shows the rest of the proteins. Subsets of proteins identified with higher aggregation propensity in naked mole-rat compared to mouse are in red and subsets of proteins identified with lower aggregation propensity are in blue. Counts of proteins from the domain analysis have lighter coloration than the counts of proteins from the whole-protein sequence analysis. Only GO terms with at least 5 proteins are shown. The groups of chaperone client proteins with significant differences of distribution compared to the rest of the proteins are marked (*).

### Supplemental Tables

All the described supplemental tables are available in the Besse_et_al_SM.xlsx file.

### Supplemental Table legends

**Table S1: Available information for Rodents from *AnAge* database**

This table contains the different metrics used for the generation of Figures 1B,C,D. Details on data collection for Maximum longevity (yrs), Body mass (g), Female maturity (days), Metabolic rate (W), and Data quality are provided in the *AnAge* database (https://genomics.senescence.info/help.html#anage).

**Table S2: Protein ortholog mapping table between naked mole-rat and mouse**

Mapping table between naked mole-rat (NKR) and mouse (M) orthologous proteins (n=13,806x2). This table was generated from after modification of the *Inparanoid* output where we selected a unique protein pair per ortholog cluster.

**Table S3: GO terms associated with domains with a significant difference in aggregation propensity**

The table was generated based on the result outputs provided by the over-representation analysis performed with the *Panther* database. It is specific to the subsets identified in the domain aggregation propensity analysis, represented by the two sections, higher aggregation propensity, and lower aggregation propensity. Within the sections, each row represents a gene ontology (GO) term associated with the following columns: information on the GO Term (GO Term, GO ID, GO Type), the number of proteins mapped to this GO term in the subset of protein used as background reference (# Reference List), the number of proteins mapped to this GO term in the analyzed subset of protein (# Analyzed List), the number of genes of expected proteins in our subset for this GO term, based on the subset of proteins used as reference (Expected), the ratio of observed number over expected number (Fold Enrichment), their associated raw P-values and FDR. +: Over-representation; -: Under-representation. Rows are sorted by descending values in the Fold Enrichment column.

**Table S4: GO terms associated to proteins with a significant difference of aggregation propensity in naked mole-rat**

The table was generated based on the result outputs provided by the over-representation analysis performed with the *Panther* database. The table is specific to the subsets identified in the per-domain aggregation propensity analysis, represented by the two sections, higher aggregation propensity, and lower aggregation propensity. The descriptions of the columns and rows are like the ones provided for Table S3.

**Table S5: Information of proteins identified in over-representation analysis with significant differences in aggregation propensity**

This table contains the proteins associated with each GO term identified from the functional enrichment analysis. The protein ID column contains the mouse Uniprot identifiers associated with naked mole-rat orthologs and that were used to retrieve their functional annotations, available in the initial FASTA file of the proteins. We also specified to which subsets (chaperone clients or other proteins) these proteins were related.
